# Robust cell tracking in epithelial tissues through identification of maximum common subgraphs

**DOI:** 10.1101/049551

**Authors:** Jochen Kursawe, Rémi Bardenet, Jeremiah J. Zartman, Ruth E. Baker, Alexander G. Fletcher

## Abstract

Tracking of cells in live-imaging microscopy videos of epithelial sheets is a powerful tool for investigating fundamental processes in embryonic development. Characterising cell growth, proliferation, intercalation and apoptosis in epithelia helps us to understand how morphogenetic processes such as tissue invagination and extension are locally regulated and controlled. Accurate cell tracking requires correctly resolving cells entering or leaving the field of view between frames, cell neighbour exchanges, cell removals and cell divisions. However, current tracking methods for epithelial sheets are not robust to large morphogenetic deformations and require significant manual interventions. Here, we present a novel algorithm for epithelial cell tracking, exploiting the graph-theoretic concept of a ‘maximum common subgraph’ to track cells between frames of a video. Our algorithm does not require the adjustment of tissue-specific parameters, and scales in sub-quadratic time with tissue size. It does not rely on precise positional information, permitting large cell movements between frames and enabling tracking in datasets acquired at low temporal resolution due to experimental constraints such as photoxicity. To demonstrate the method, we perform tracking on the *Drosophila* embryonic epidermis and compare cell-cell rearrangements to previous studies in other tissues. Our implementation is open source and generally applicable to epithelial tissues.

## 1 Introduction

Live-imaging microscopy is a powerful, and increasingly quantitative, tool for gaining insight into fundamental processes during embryonic development [1–3]. Quantitative information on cell growth, proliferation, death, shape changes and movement extracted from live-imaging reveals how such processes are regulated to give correct tissue-level behaviour. This approach has been particularly successful in characterising the growth and patterning of embryonic epithelial tissues in a number of model organisms [4–9].

A common experimental technique for visualising cell shapes in an epithelial sheet is to fluorescently tag a molecule marking cell boundaries, such as E-cadherin (figure 1A). The analysis of time-lapse microscopy data obtained from such tissues is extremely challenging [2, 3], especially in cases of imaging data of rapidly evolving tissues, and when limitations of, for example, microscope speed, imaging resolution or phototoxicity prohibit the creation of datasets with high temporal and spatial resolution.

**Figure 1:**
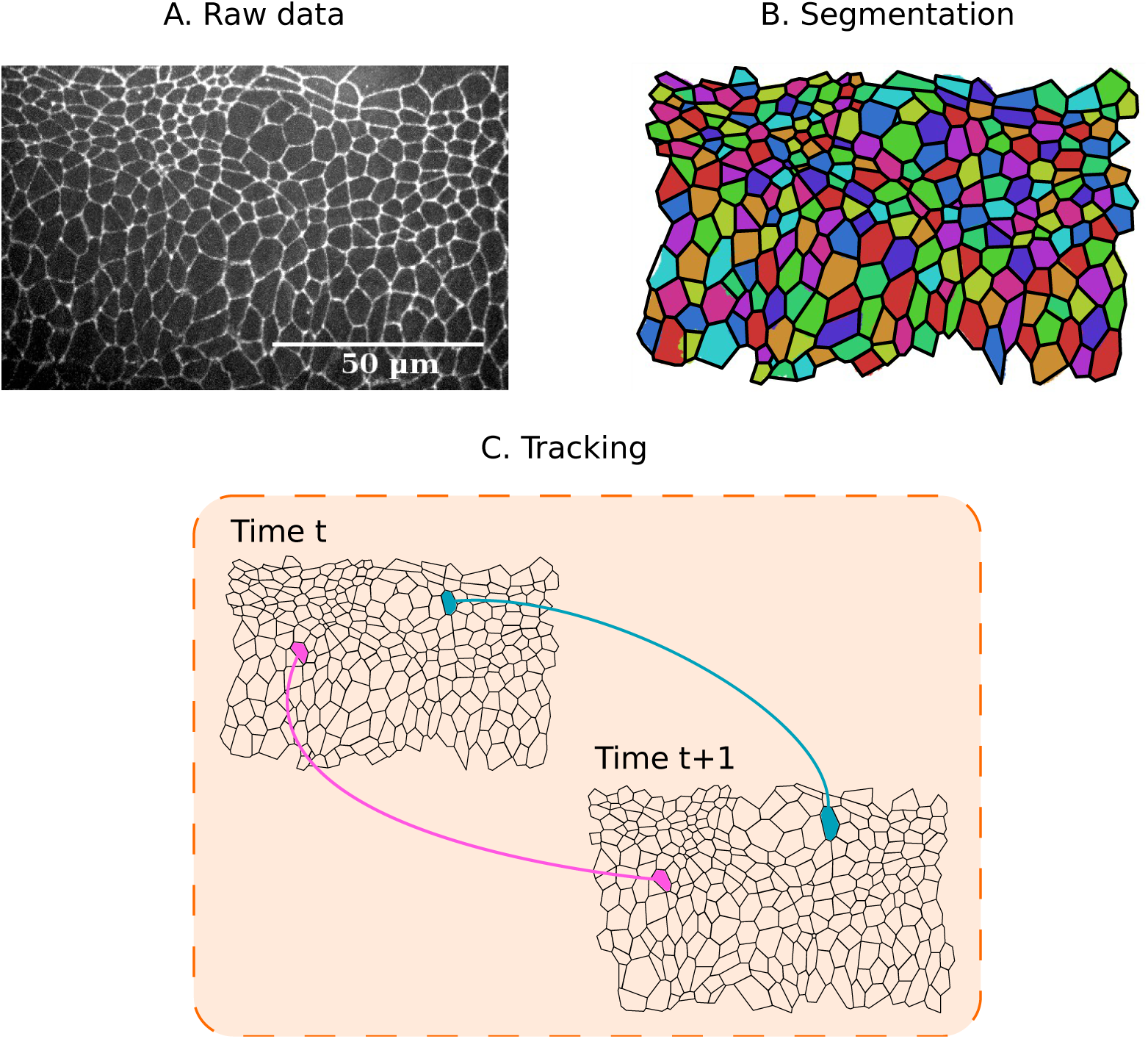
Pipeline for analysing epithelial tissues. (A) Example raw data. Frame of a live-imaging microscopy video of the lateral epidermis of a stage-11 *Drosophila* embryo, expressing DE-Cadherin::GFP. See Experimental Methods for details. (B) Segmentation of this image, showing cell shapes (coloured regions) and polygonal approximation based on three-cell junctions (black lines). See Methods section for details of segmentation. (C) Cell tracking involves registering individual cells across consecutive segmented images.

The analysis of time-lapse microscopy data comprises two major steps: segmentation and tracking (registration). Segmentation must be performed for each frame of a video and involves the identification of objects and landmarks, such as cell shapes (figure 1B). Automated segmentation is hindered by various factors such as noise in fluorescent signals, uneven illumination of the sample, or overlapping cells in a two-dimensional projection. Often, manual correction is necessary to address over-segmentation, where too many cells are detected, or under-segmentation, where too few cells are detected [10–12]. Tracking involves the association of segmented cells across video frames (figure 1C) and requires resolving cellular movement, cell division, cell death, and cells entering and leaving the field of view [12].

Numerous algorithms are available for the segmentation and tracking of cellular-resolution microscopy data [10, 11, 13]. Common methods for cell tracking utilize optimization techniques to minimise differences in cellular properties between two frames [11, 14–17]. The min-cost max-flow algorithm [14] uses linear integer programming to minimise differences in cell areas, perimeters, orientations, and locations between frames, whereas multiple-parameter tracking [15] employs global optimization to minimize differences in cell shapes as well as locations. In contrast, multitemporal association tracking [16, 17] minimises differences in cell locations and sizes by using a probabilistic approach that finds the most likely extension to existing cell trajectories. Chain-graph models [18] minimise differences in cell velocity while overcoming missegmentation by verifying that each segmented object continues or begins a cell trajectory in successive frames. Optical flow (‘warping’) between successive frames can be used to guide cell tracking as well as segmentation [19]. It is also possible to combine segmentation and tracking of 2D microscopy videos by interpreting time as a third spatial dimension and employing 3D segmentation techniques [20]. The nearest-neighbour method associates two cells in consecutive frames with each other if their respective centroids have minimal distance within the field of view [10], or if their overlap in pixels within the field of view is maximal [21, 22]. Particle image velocimetry, a technique originally developed to analyse fluid flow [23], has also been employed to track cells in epithelial tissues [24].

Software implementations and computational tools for cell tracking include FARSIGHT [25] (segmentation only), SeedWaterSegmenter [10] (nearest-neighbour tracking), ilastik [18] (chain-graph models), Tufts Tissue Tracker [11] (min-cost max-flow algorithm), Tracking with Gaussian Mixture Models [26] (nearest-neighbour tracking), Packing Analyzer [27] (particle image velocimetry) and EpiTools [13] (nearest-neighbour tracking). These algorithms and software tools primarily rely on there being small differences in cell positions and shapes across consecutive images. Their performance is therefore hindered when analysing data from *in vivo* studies where phototoxicity provides a barrier to high temporal resolution imaging [28–30]. To address this limitation, we propose a novel algorithm for cell tracking that uses only the connectivity of cell apical surfaces (figure 1). By representing the cell sheet as a physical network in which each pair of adjacent cells shares an edge, we show that cells can be tracked between successive frames by finding the *maximum common subgraph* (MCS) of the two networks: the largest network of connected cells that is contained in these two consecutive frames. It is then possible to track any remaining cells based on their adjacency to cells tracked using the MCS. Our algorithm does not require the tuning of parameters to a specific application, and scales in subquadratic time with the number of cells in the sheet, making it amenable to the analysis of large tissues.

We demonstrate here that our algorithm resolves tissue movements, cell neighbour exchanges, cell division, and cell removal (for example, by delamination, extrusion, or death) in a large number of *in silico* datasets, and successfully tracks cells across sample segmented frames from *in vivo* microscopy data of a stage-11 *Drosophila* embryo. We further show how our algorithm may be used to gain insight into tissue homeostasis by measuring, for example, the rate of cell rearrangement in the tissue. In particular, we find a large amount of cell rearrangement within the observed dataset despite the absence of gross morphogenetic movement. The remainder of the paper is structured as follows. In Section 2 we describe the algorithm for cell tracking. In Section 3 we analyse the performance of the algorithm on *in silico* and *in vivo* datasets. Finally, in Section 4 we discuss future extensions and potential applications.

## 2 Methods

In this section we provide a conceptual overview of the core principles underlying our cell tracking algorithm. We focus on providing an accessible, non-technical description rather than including all details required to implement the algorithm from scratch. A comprehensive mathematical description of the algorithm is provided in Supplementary Material S1.

The input to the algorithm is a set of segmented images obtained from a live-imaging microscopy dataset of the apical surface of an epithelial cell sheet. For each image, the segmentation is assumed to have correctly identified which cells are adjacent and the locations of junctions where three or more cells meet. Various publicly available segmentation tools can be used for this segmentation step, for example SeedWaterSegmenter [10] or ilastik [18]. The segmentation is used to generate a polygonal approximation to the cell tessellation (figure 1B-C). Such approximations are an adequate assumption for many epithelia [11, 31–34].

Our algorithm tracks cells by interpreting the polygonal representations arising from the segmentation as networks (‘graphs’) of cells. Examples of such networks are shown in figure 2A. In this representation, each cell corresponds to a vertex of the network, and two vertices are connected by an edge if the corresponding cells are adjacent. Our algorithm tracks cells across consecutive images by aligning the networks of cells that correspond to these images. This network alignment is achieved in three steps. First, we generate an initial tracking for subsets of the cells in each pair of consecutive images by finding the MCS between the two corresponding networks (figure 2B). Second, this MCS is reduced to avoid tracking errors (figure 2C). Third, remaining untracked cells are tracked based on their adjacency to cells within the MCS (figure 2D). In the final output of the algorithm, each tracked cell of a frame is paired with exactly one cell in the subsequent frame.

**Figure 2:**
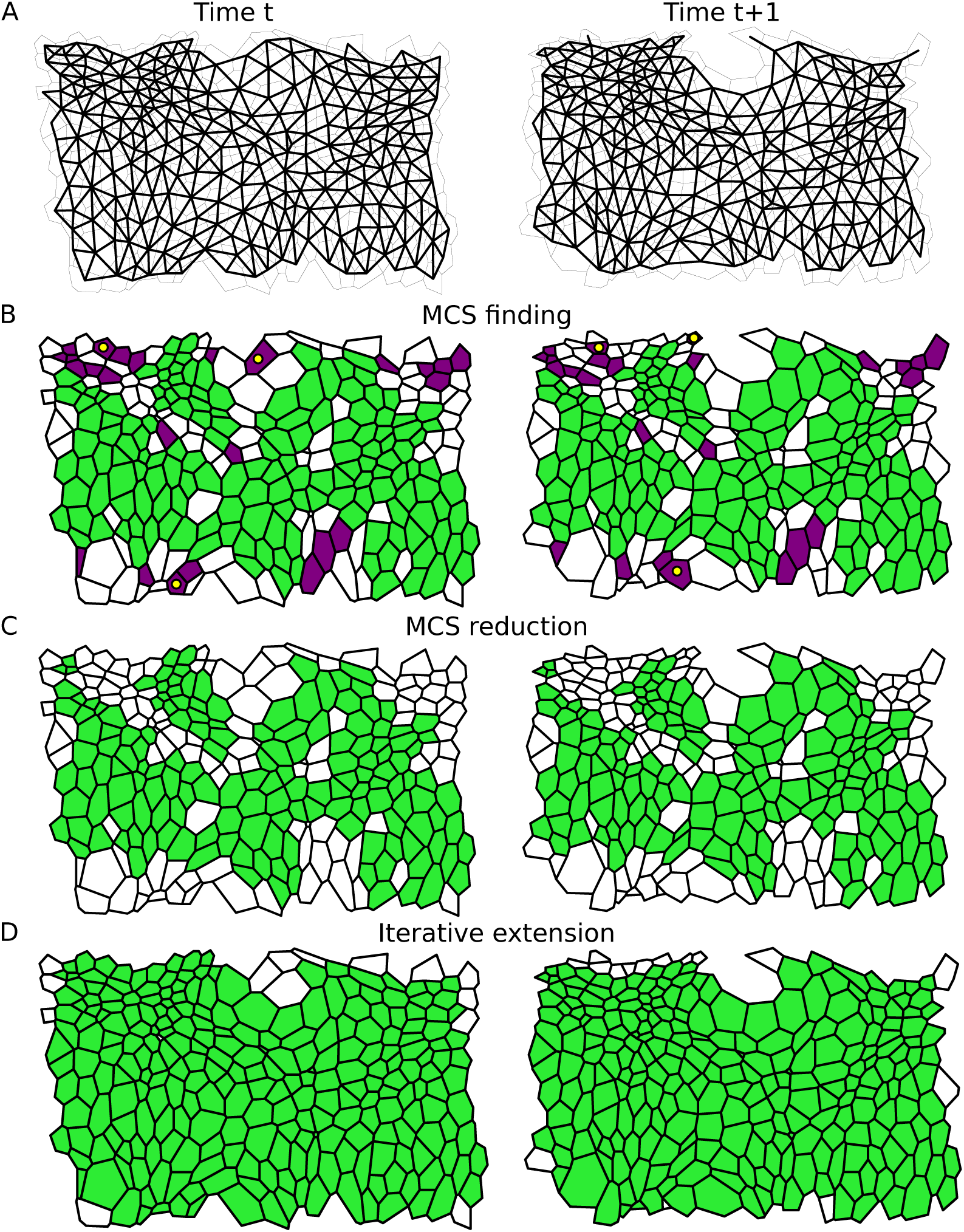
Illustration of our cell tracking algorithm. (A) Grey: Two consecutive segmented time-lapse images (left and right columns) of the lateral epidermis of a stage-11 *Drosophila* embryo, taken five minutes apart. See Experimental Methods for details. There are several cell neighbour exchanges between these images. Black: Overlay of the network of cells that the algorithm uses for cell tracking. Cells in the tesselation correspond to network vertices that are connected by an edge if the cells are adjacent. (B) We first identify a cell mapping between the two graphs based on the MCS. This includes correctly tracked (green/light) cells and cells that have only few tracked neighbours (purple/dark). Here, the MCS incorrectly tracks three cells (yellow/light dots). (C) Weakly connected cells and small isolated clusters of cells are removed from the MCS to prevent mismatches. (D) An extended tracking mapping is constructed, which includes the maximum possible number of cells. See Methods section for details. The remaining white cells have entered or left frame of view between images and therefore are not tracked.

The key step in this network alignment approach is the identification of a MCS [35, 36]. A MCS comprises the largest sub-network that is contained in two larger networks; thus finding an MCS can be understood as recognising patterns of connections that are preserved between two networks. In the present work, the structure of the MCS roughly corresponds to cells that do not rearrange between consecutive images, except for few cells at its boundaries.

In figure 2B, we visualize the MCS generated by our algorithm as a collection of green (light) and purple (dark) cells. Most of the highlighted cells in figure 2B are tracked correctly by the MCS. Three cells in each frame are marked by a yellow (bright) dot. Within the two cell networks, these cells are members of the MCS. However, these cells are not tracked correctly by the MCS. This mismatch arises since the MCS is found based on the connectivity of cells within the network alone. The fewer connections a cell has to other cells in the MCS, the less information about the cell’s position and shape is encoded by these network connections, and so the greater the possibility of mismatches. To avoid such tracking errors, we remove any cells that have only a few connections within the MCS, as well as small isolated clusters of cells. All cells that are removed from the tracking in the second step of our algorithm are shown in purple (dark) in figure 2B. In figure 2C we highlight the cells that are tracked after applying this second step of our algorithm.

Cells that are untracked after reducing the MCS are then tracked based on their connections to previously tracked cells. This last step of our algorithm comprises starting from the MCS and iteratively ‘growing’ the set of tracked cells by adding cell-to-cell matches to the tracking that maximise the number of preserved connections to other tracked cells. In this step, the algorithm also resolves cell neighbour exchanges, cell removals, and cell divisions by identifying changes in the network structure that are characteristic of these events. For example, a cell neighbour exchange corresponds to the deletion of a network connection while a new connection is added.

### The MCS is identified through repeated seeding and iterative extension

In computer science, MCS finding has been known to be an NP-hard problem [35, 36]: the time to find an exact MCS of two networks increases exponentially with the size of the networks, which poses a computational barrier to the use of MCS-finding algorithms in applications. We overcome this computational barrier in the present work by constructing the MCS iteratively from the MCSs of smaller subgraphs, exploiting the planar structure of our cell networks to reduce the complexity of the problem.

To start the construction of the MCS, the algorithm identifies a match between two cells in the consecutive images for which the structure of the network of their surrounding cells is identical. Here, the network of surrounding cells is restricted to the network formed by a cell’s neighbours and its second nearest neighbours (see figure 3B). If no such initial match can be found, the algorithm instead searches for an initial match where only the first-order neighbourhood is preserved, under the condition that this neighbourhood does not touch the boundary of the tissue. This latter condition avoids tracking errors that can occur on the tissue boundary where cells have few neighbours.

**Figure 3:**
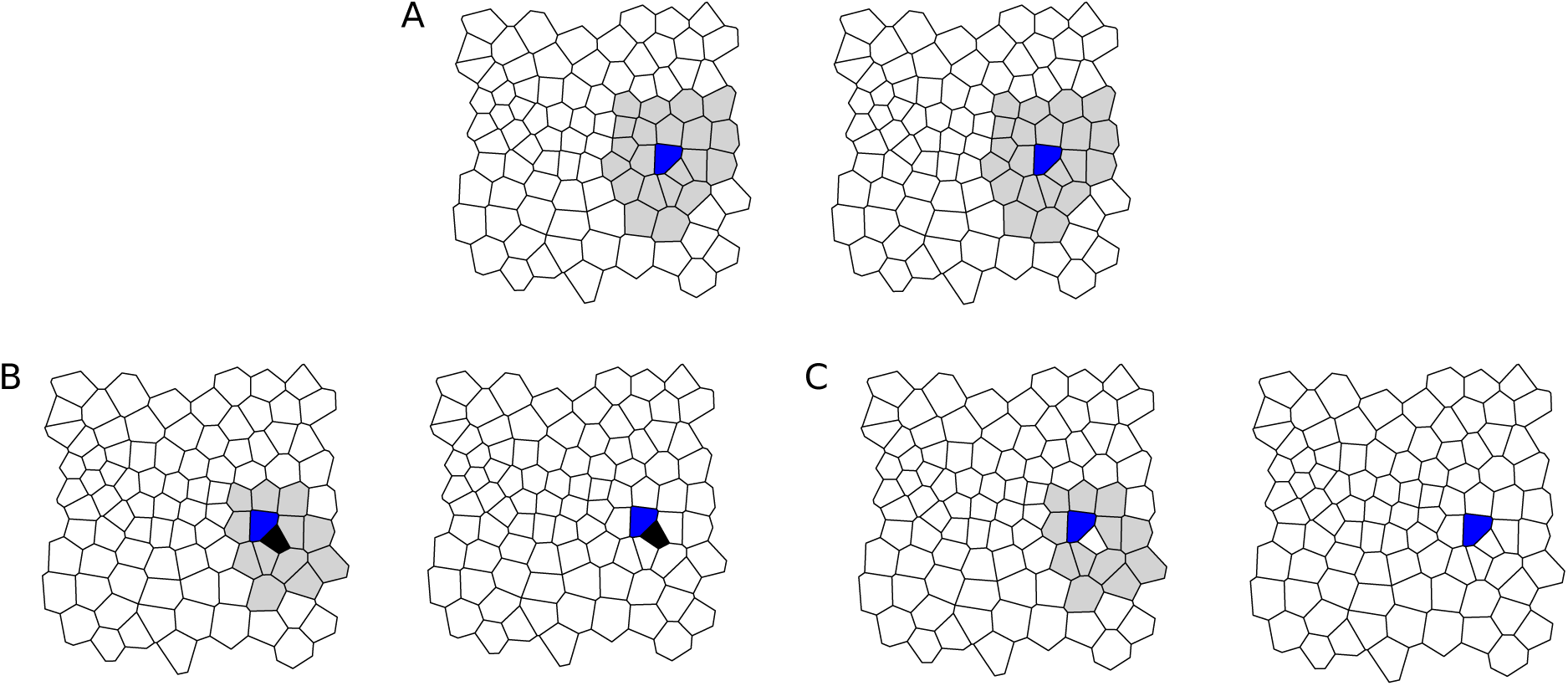
Construction of the MCS. (A) The algorithm picks a first match of cells for the MCS (blue) if their neighbourhoods form identical networks. The considered neighbourhood (grey) includes all neighbours and second nearest neighbours. (B-C) Additional cells are added to the MCS iteratively by inspecting the MCS between the grey area on the left, and the white area on the right. In (B), where the black cell is paired correctly, the local MCS is larger than in (C), where the selected cell is not considered for mapping. Hence, the pairing of black cells is added to the MCS.

Once the initial match (a ‘seed’) is found, the algorithm iteratively adds further cells to the MCS. At each step of this iteration, a cell in the first network is picked that is adjacent to the existing MCS and has a minimal number of potential matches. This number of potential matches is determined based on how many cells in the second network have the same number of neighbours as the considered cell while preserving connections to already tracked cells. Among the choice of potential matches the algorithm identifies an optimal match based on the local network structure of these cells’ neighbours. A cell in the second network is identified as an optimal match if the network structure of its neighbourhood is most similar to the cell in the first match. This choice is made based on local MCSs between the neighbourhood-networks of the cell in the first image and each potential match. Note that the optimal choice may exclude the cell from the tracking entirely. In this case, most neighbours are included in the local MCS when the considered cell is not tracked, indicating for example a cell removal event. In this case the cell is not mapped and the algorithm proceeds by inspecting another cell in the first match. Cells in the first frame for which no match in the second frame has been found may be re-inspected at later stages of the algorithm as the size of the identified MCS increases. Once no more adjacent cells can be added to the MCS through this iterative extension, the iteration continues the search among untracked cells in the first network that are not adjacent to the existing MCS. As soon as at least one cell has been added to the MCS in this way, the algorithm again restricts its search to adjacent cells. The algorithm halts once no further cell-to-cell matches can be found. During the construction of the MCS the algorithm ignores any potential cell-to-cell matches where the corresponding cell centroids are more than a cutoff distance *d*_max_ apart within the field of view. Throughout the manuscript, we choose *d*_max_ to be ten average cell lengths.

Once the MCS is complete, any cells that have less than three isolated connections to other cells in the MCS are removed from the tracking. Any clusters of ten or fewer cells are also removed from the tracking result. Both of these steps help to minimise tracking errors (figure 2B-C).

### Cells are added to the tracking result by inspecting connections to previously tracked neighbours

Through the identification of the MCS the algorithm tracks most of the cells that do not rearrange between consecutive frames. Next, the algorithm tracks any remaining cells, and identifies cell rearrangements, cell removal, and cell division events. Similar to the construction of the MCS, the tracking of remaining cells is iterative. At each iteration, the algorithm identifies a cell-to-cell match that maximises the number of connections to already tracked cells, thus ‘growing’ the set of tracked cells from the intermediate tracking result of the MCS. When adding cells to the tracking, the algorithm ensures that a cell cannot gain more tracked neighbours between consecutive frames than the number of tracked neighbours preserved between these frames. The algorithm also requires a cell to have at least two tracked neighbours in order to be added to the tracking in this way.

Once all possible cells have been tracked, the algorithm resolves division events. Division locations can be identified as regions in the second frame that contain more cells than the corresponding region in the first frame. Since the algorithm will have found exactly one match in the second network for each tracked cell in the first network and *vice versa*, there are thus untracked cells in the second frame wherever a cell divides between two consecutive frames. The algorithm attempts to resolve divisions events by identifying changes in cell-to-cell connectivity that are characteristic to dividing cells (figure 4). For example, two cells adjacent to each division must gain a neighbour (grey cells in figure 4A), and in many cases the mother and daughter cells are easily identified as the cells that are shared neighbours of these cells adjacent to the division event. However, one of the daughter cells may be four- or three-sided (figure 4B-C). In these cases, the algorithm is not able to determine the mother- and daughter cells based on their network properties alone. Instead, the algorithm takes the geometric shape of the cells into account. The mother and daughter cells are chosen by identifying which pair of potential daughter cells has the closest position to their potential mother cell.

**Figure 4:**
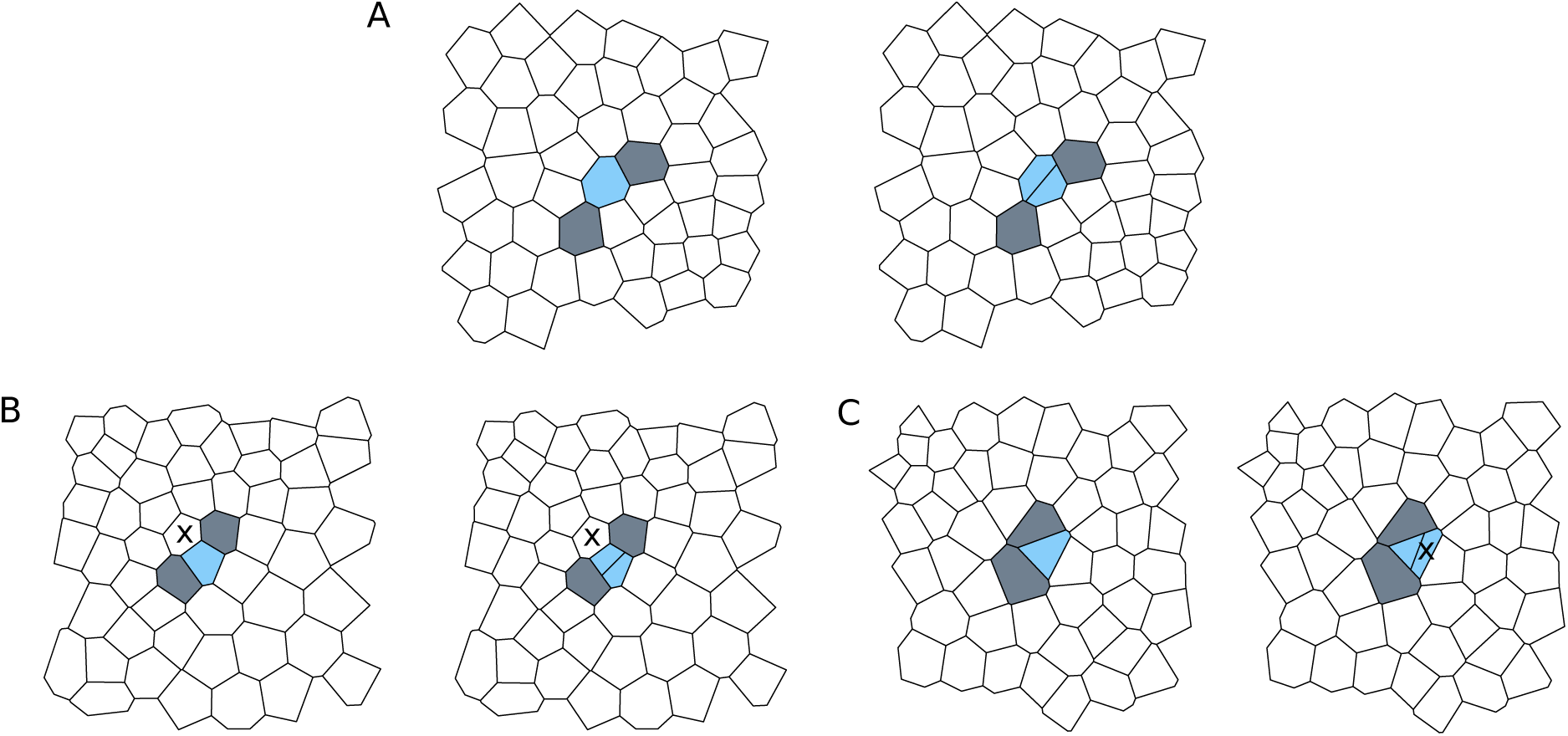
Resolving division events. Dividing cells are coloured blue. (A) Division events are resolved by identifying cells that gain an edge between the time frames (grey). The dividing cell and the daughter cells are shared neighbours of such cells. (B) When one of the daughter cells is four-sided, two mother cells are possible, the blue marked mother cell, and the cell marked by an ‘x’. (C) If one of the daughter cells is three-sided, the mother cell can be mistaken for having gained an edge if it is identified with the daughter cell labelled ‘x’. Our algorithm correctly resolves each of the types of division events shown in (A)-(C).

Cell deaths are identified as cells in the first frame that do not have a tracked match in the second frame and that are not on the boundary of the region of tracked cells.

### Code availability

The code used in this article is publicly available under the 3-clause BSD license as the MCSTracker project (https://github.com/kursawe/MCSTracker). The project is implemented in pure Python, employs unit testing [37] and is fully documented. Graphs in our code are represented using the Python package NetworkX [38].

### Generation of *in silico* datasets

To test the algorithm, we generate *in silico* datasets that include examples of cell divisions, removals and neighbour exchanges, as well as tissue movement. These datasets are generated using Voronoi tessellations modified using Lloyd’s relaxation, which resemble cell packings in a variety of epithelial tissues [33, 39].

To generate polygonal patterns of size *m × n*, where *m* and *n* are natural numbers, (*m* + *g*) *×* (*n* + *g*) Voronoi seeds are distributed uniformly at random in a 2D domain Ω of width *m* + *g* and height *n* + *g* (figure 5A). Here, *g* denotes the size of a boundary region that is introduced to reduce the impact of the Voronoi boundary on the patterns. The domain Ω is surrounded by two additional rows of evenly spaced seeds on each side. The inner row is a distance of 0.5 length units to Ω, and the seed-spacing is 1.0. The outer row has a distance of 1.5 to Ω, and the seeds are shifted parallel to the first row by a distance of 0.5. The Voronoi tessellation of all these seeds is then constructed.

**Figure 5:**
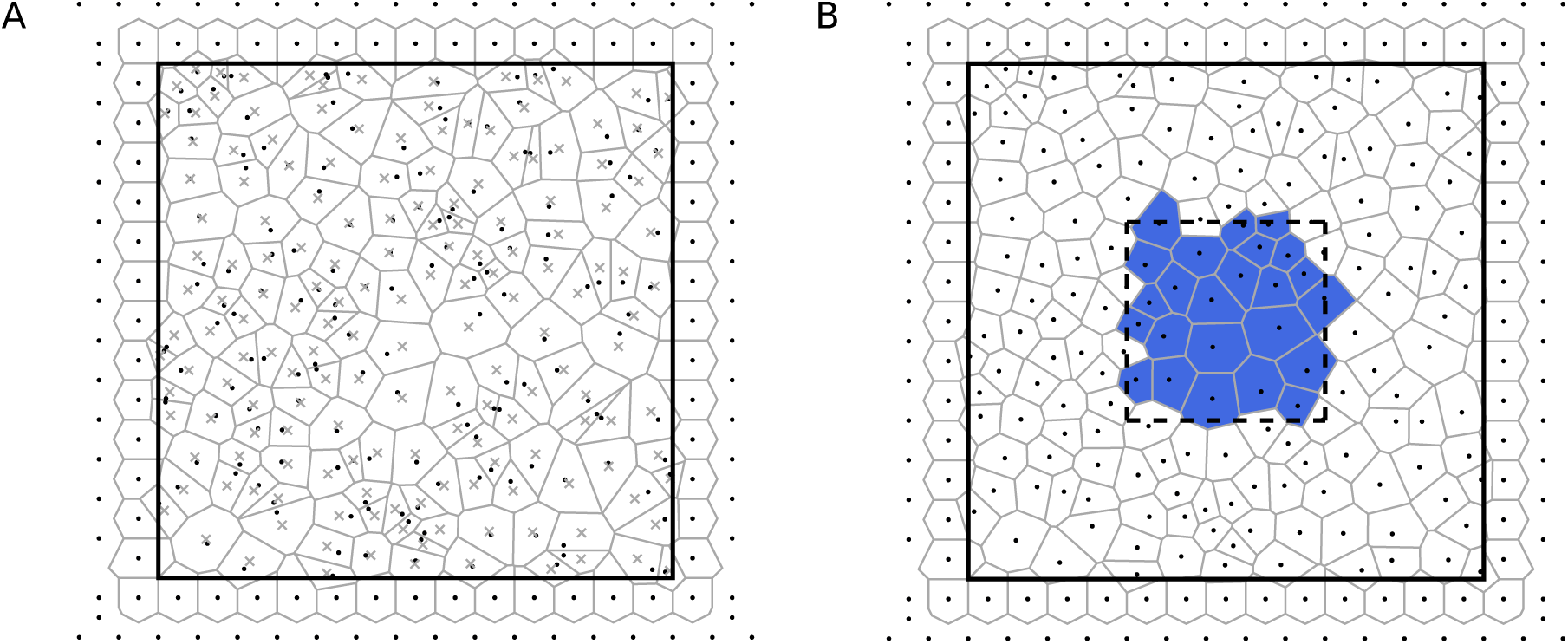
Generation of *in silico* data. (A) Random seeds (black dots) are placed inside a domain Ω (black line). Additional seeds are placed outside Ω. The Voronoi tessellation of all seeds is shown in grey, excluding Voronoi regions corresponding to the outermost row of seeds, since these are large or unbounded. The centroids of the Voronoi regions (grey crosses) differ from the seeds. (B) The centroids of the Voronoi regions in (A) are used as seeds for a new Voronoi tessellation, for which evenly spaced seeds are again added outside the domain Ω. Voronoi regions whose centroids lie within a central rectangle (dashed black line) are collected to form the *in silico* tissue (blue). In this figure, one Lloyd’s relaxation step is shown. Throughout this study, we generate *in silico* tissues using four Lloyd’s relaxation steps.

In each Lloyd’s relaxation step, the polygons (or infinitely large areas) corresponding to the regularly spaced seeds outside Ω are removed from the tessellation. Next, the centroid of each remaining polygon is calculated and registered as a new seed. Further seeds are added that again correspond to two rows of evenly spaced seeds outside Ω. A new Voronoi tessellation is then constructed (figure 5B). This procedure is repeated for *L* relaxation steps, after which all generated polygons are discarded except those whose centroids lie within a rectangular domain of size *n × m* area units whose centroid coincides with that of Ω (figure 5B).

The polygonal tessellations have approximately *m × n* polygons of average area 1.0. During the generation of the tessellations, evenly spaced seeds outside Ω are added to prevent the occurrence of infinitely large polygons inside Ω. The boundary of size *g* is added in between the generated tessellation and the evenly spaced seeds to reduce the effect of the evenly spaced boundary seeds on the tessellation. Throughout this study, we use *g* = 8 and *n_L_* = 4, resulting in cell packings similar to those observed, for example, in the *Drosophila* wing imaginal disc [33]. We provide further details of how tissue rearrangements are implemented in the Results section.

### Experimental methods

Live-imaging of cell proliferation was performed in stage-11 *Drosophila* embryos expressing a tagged version of DE-Cadherin (DE-Cadherin::GFP) using a spinning disc confocal microscope, as described in [40]. For the embryo setup, a modified version of the standard live-imaging protocol was used [41].

### Data segmentation

Microscopy images were segmented using pixel classification in ilastik [18]. The classifier was trained to recognise cell outlines and the segmentation of each frame was manually corrected. A watershed algorithm was used to identify the precise shape of the cell outlines. Each segmented frame was converted to a 16-bit grayscale image where pixels belonging to different cells had different integer values. Polygonal tessellations for the tracking algorithm were generated from the segmented image in two steps. First, all junctions between three or more cells were identified as points where pixels of three or more different cells met; second, vertices were assigned to cells. Then, edges shorter than two pixels (0.5 µm) were removed and replaced by a single vertex at the midpoint of the edge. Finally, polygons at the boundary of the tissue were removed from the simulation. This removal was necessary since cell shapes at the tissue boundary are poorly approximated by polygons due to missing vertices. Note that our algorithm can interpret segmentations saved using either ilastik [18] or SeedWaterSegmenter [10].

## 3 Results

### *In silico* testing of the algorithm

To assess the performance of the algorithm, we begin by applying it to *in silico* datasets that include cell neighbour exchanges, tissue movement, cell removal and cell division, respectively. In each case, we compare the outcome of the tracking algorithm to the ground truth.

We begin by assessing the ability of the algorithm to resolve permutations in otherwise identical tissues (figure 6A). In this test, a random tessellation of size nine by nine cells is created as described in the Methods section, and integer identifiers *c*_*i*_ are assigned to each cell. Next, an identical copy of the tissue is created in which the integer identifiers are randomly shuffled. A ground truth mapping from the first to the second integer identifiers is generated. Next, the algorithm is applied. Upon conducting 100 such tests, we find that all identified cell-to-cell mappings are matched correctly, as compared to the ground truth. In rare examples, isolated cells at the boundary of the tissue are are not tracked. In these examples, either a single cell has only one adjacent cell in the tissue, or two cells of identical polygon number are adjacent and share exactly one neighbour. Neither the MCS detection algorithm, nor the post-processing algorithm are able to resolve such mappings, which involve fewer than four cells in each dataset (fewer than five percent of the tissue).

**Figure 6:**
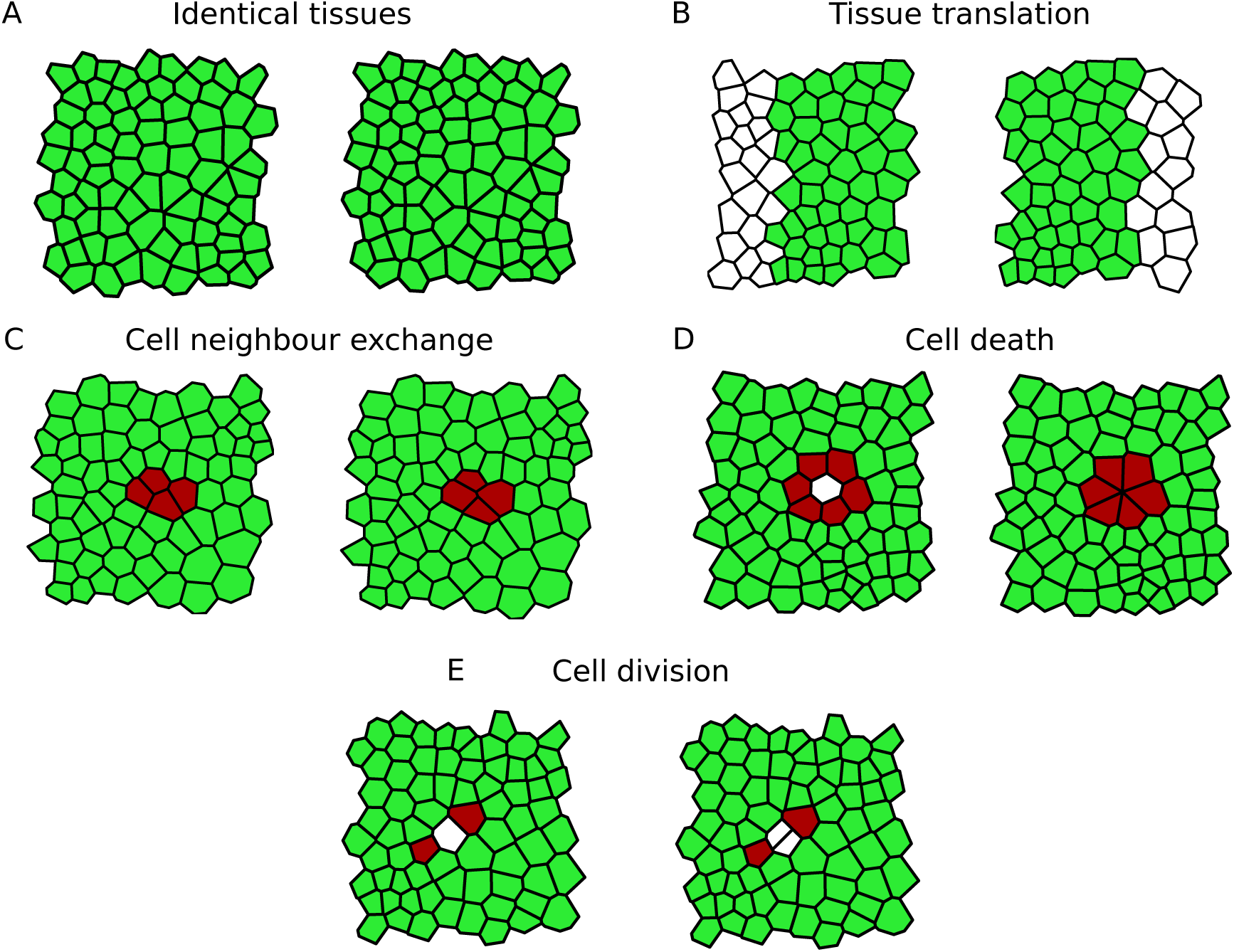
Examples of *in silico* test cases. In each image, cells identified by the MCS algorithm are highlighted in green (light), whereas cells that have been filled in by the post-processing steps are highlighted in red (dark). The algorithm tracks cells between identical tissues (A), in tissues undergoing translation (B), cell neighbour exchange (T1 transition) (C), cell removal

We design four further tests of tissue rearrangements (figure 6B-E). The first test comprises tissue movements between images (figure 6B). In this test, a tissue of size fifteen by eight cells is generated as described in the Methods section. Two smaller tissues of width seven units are cut out of this tissue, which each cover the full height of the tissue, and which are horizontally translated relative to each other by a distance of two cell lengths. The position of each three-cell junction in both tissues is shifted such that the *x*-coordinate of the left-most junction in each tissue is 0.

The second test (figure 6C) generates cell neighbour exchanges, also called T1 transitions [42, 43]. In our implementation of T1 transitions, an edge shared by two cells is replaced by a new perpendicular edge (of length *l*_T1_ = 0.2 units) such that the local cell connectivity changes (figure 2B). We create two identical copies of a tissue of size nine by nine cells. In the second copy, a T1 transition is performed on an edge in the centre of the tissue.

The third test involves cell removal (figure 6D). In this test, we first generate two identical copies of a tissue of size nine by nine cells. In the second copy, we replace the central cell by a vertex shared by its neighbouring cells, a rearrangement similar to so-called T2 transitions [42]. The final test involves cell divisions (figure 6E). Here, we once again create two identical copies of size nine by nine cells. In the second copy, a cell in the centre of the tissue is bisected by introducing a straight line in a random direction through the centroid of that cell.

For all tests generated in this way, integer cell identifiers in the second tissue are randomly shuffled, and a ground truth is generated. We run 100 realisations of each test case, and compare the tracking outcome to the ground truth. In all cases we find that cells are tracked correctly, with at most three unmatched cells at the boundary of the sheet.

In figure 6, all cells identified after the cleaning step, in which weakly connected cells are removed from the MCS, are coloured green, whereas cells that are identified by the post-processing algorithm are coloured red. Note that the exact number of cells that are identified by the post-processing algorithm varies between individual realisations of the tests. In many cases, the cells identified by the post-processing algorithm include cells that are adjacent to those undergoing division, removal or neighbour exchange.

We next analyse the extent to which the success of our tracking algorithm depends on the number of Lloyd’s relaxation steps, *n*_*L*_, used to generate the *in silico* datasets. To investigate this we iteratively increase *n*_*L*_, thus generating tissues with increasingly homogeneous graph structures, and repeat all tests. We find that the algorithm successfully passes all tests for all values of *n*_*L*_ from 4 up to 14.

### Algorithm performance for large numbers of cell neighbour exchanges

To assess the performance of the algorithm when applied to tissues exhibiting large numbers of cell neighbour exchanges, we next apply the algorithm to *in silico* datasets with increasing numbers of cell neighbour exchanges between frames (figure 7). The number of correctly tracked cells decreases as the number of cell neighbour exchanges increases. However, the number of incorrectly tracked cells remains below 20% throughout the analysed range of neighbour exchanges, and decreases to zero as the number of edge swaps exceeds 10%.

**Figure 7:**
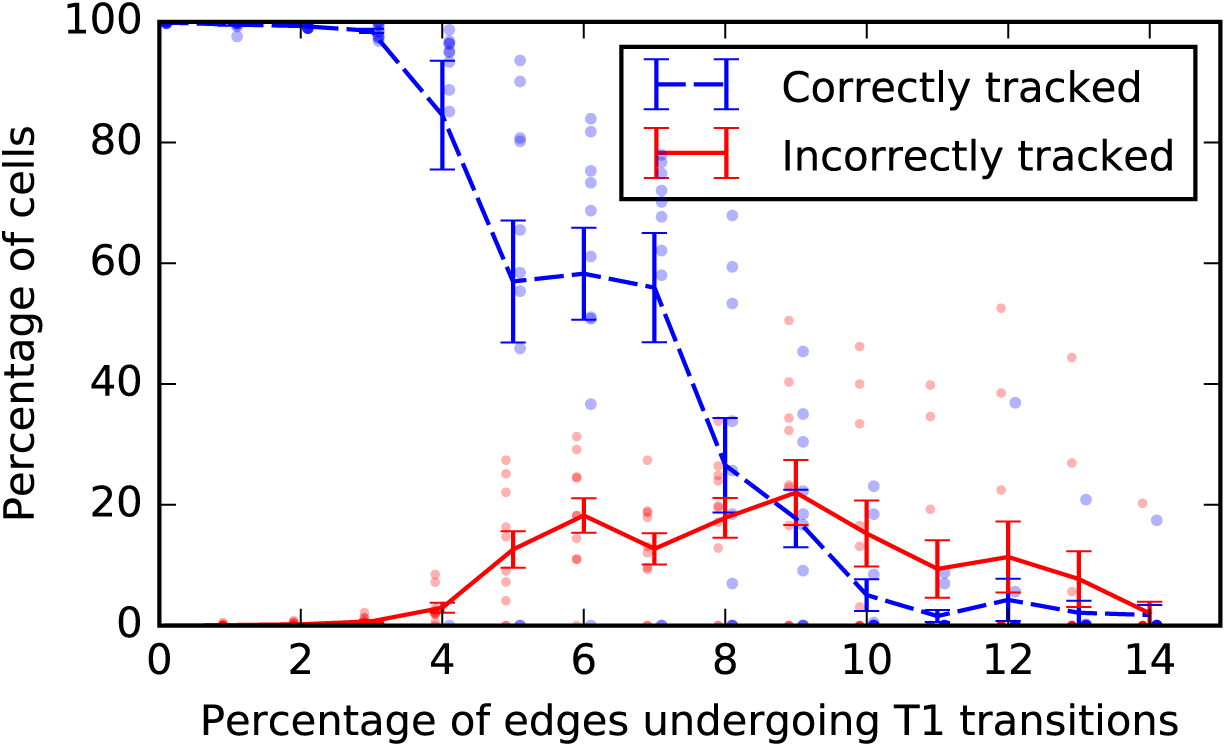
Success rate of the algorithm for *in silico* tissues with increasing frequency of cell rearrangement. Virtual tissues spanning 20 cell lengths in each dimension are generated, and T1 transitions are applied to an increasing proportion of the inner edges of the tissue. For each ratio of T1 transitions, 10 repetitions of the test are run, and the ratio of correctly and incorrectly tracked cells in the tissue is recorded. The dashed blue and solid red lines correspond to mean values of correctly and incorrectly tracked cells, respectively. Error bars denote the standard deviation of the mean, and results of individual runs of the test are represented by dots. When 3% of the edges in the tissue undergo T1 transitions, roughly 25% of the cells exchange neighbours.

The number of untracked cells increases rapidly as the percentage of cell-cell interfaces that are swapped between successive images increases from five to ten percent. Note that the percentage of cells involved in these neighbour exchanges is larger than the percentage of cell-cell interfaces that are swapped, since an individual T1 transition changes the cell neighbour relations of four cells, and each cell shares multiple inner edges. For example, rearranging five percent of the inner edges of the tissue affects roughly 40% of the cells in the tissue, while rearranging ten percent of the tissue edges affects up to 70% of the cells. The number of (correctly or incorrectly) tracked cells drops to zero if the tissue rearranges so much that the neighbourhood of each cell changes; in this case a first match cannot be found to initialise the MCS construction algorithm.

### Application of the algorithm to *in vivo* data

Figure 8 shows the first three of 21 segmented image frames of the lateral epidermis of a stage-11 *Drosophila* embryo to which the algorithm was applied. During stage 11, gross morphogenetic movements do not occur but the tissue is very active with a large number of proliferations occurring within a short duration, making this a much more challenging tissue on which to perform cell segmentation and tracking than the wing imaginal disc, where many previous efforts have been made [4, 13, 31, 44]. Cell delamination is also more common than in the wing imaginal disc during normal development. This stage of development thus offers a true test of the present algorithm.

**Figure 8:**
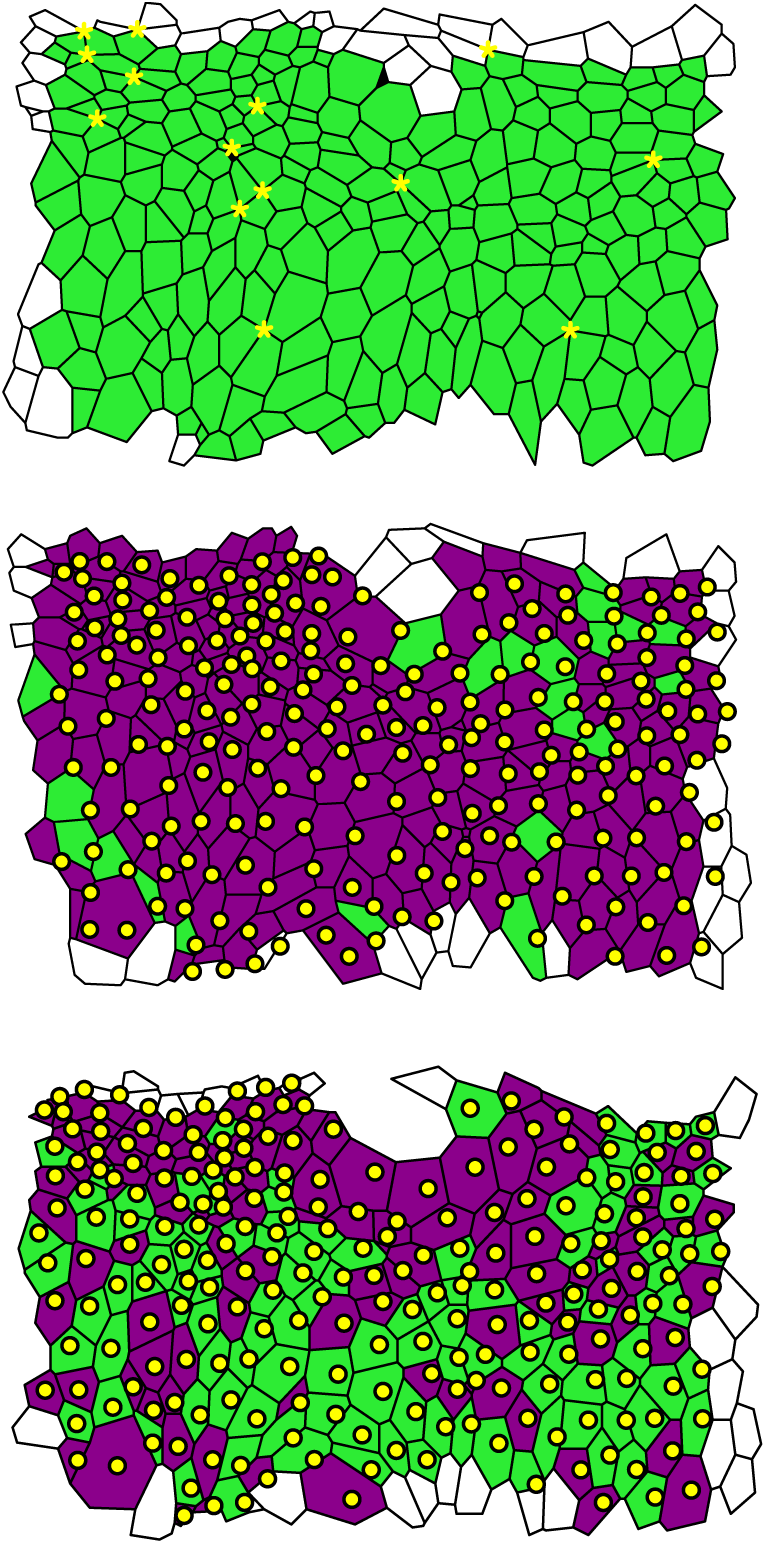
Three segmented data frames of an *in vivo* time-lapse microscopy video of the lateral epidermis of a stage-11 *Drosophila* embryo. Cells that are tracked across all frames are coloured green or purple, and cells that leave or enter the tissue at the boundary are white. Dying cells are black. The centroids of tracked cells of the respective previous frames are included as yellow dots, and cells that contain only their centroid from the previous frame are coloured green, whereas cells that do not contain their centroid from the previous frame, and cells that contain multiple centroids, are coloured purple. Together, the centroids and the colouring illustrate that it is challenging to track cells between the data frames using solely centroid positions. Yellow asterisks in the first frame denote higher-order junctions where more than three cells meet.

The images were taken five minutes apart over a time span of 100 minutes. These first three images comprise 271, 263 and 263 cells, respectively. Our algorithm tracks 247 cells between the first and second images, 245 cells between the second and third images and 234 cells across all three images. The centroids of cells of previous images are superimposed on the tracking results in figure 8, illustrating that the algorithm successfully tracks cells in situations where it is difficult to match cells between images based on the centroid positions alone. Cells that include only their corresponding centroid from the previous image are coloured in green, while cells that do not include their corresponding centroid from the previous image, and cells that include multiple centroids from the previous image, are coloured in purple. In the first frame we highlight ‘higher-order’ junctions (shared by four or more cells) by yellow asterisks. Such junctions occur frequently throughout the dataset.

On average, cell centroids move 0.75 cell lengths between the first and second images, with a maximal displacement of 1.17 cell lengths. Between the first and second images 36 cells undergo a net gain in edges, whereas 20 cells have a net loss of edges. In total, four cell deaths and no cell divisions are observed across the three data images. Inspection of all individual cell tracks reveals that none of the cells are tracked incorrectly.

The data in figure 8 are the first three out of 21 frames. In figure 9 we show the results of the analysis of the full dataset, including all 21 frames. During the period of measurement the total number of cells increases from 280 to 330 cells, whereas the total number of tracked cells increases from 270 to roughly 310 cells. As the number of cells in the tissue rises, the total number of cell rearrangements increases, whereas the average cell area decreases. Here, the number of cell rearrangements is measured by counting how many cells change their cell-neighbour number between consecutive frames. For all frames, the number of tracked cells is lower than the number of cells in the tissue. A visual inspection of the tracked data reveals that the difference between the total number of cells and the number of tracked cells is largely due to cells entering or leaving the field of view. The percentage of cells that our algorithm tracks is lowest (84%) when the rates of cell division and cell rearrangement are highest, which occurs at 70 minutes. Here, the number of tracked cells decreases since the algorithm is not yet able to resolve division events immediately adjacent to rearrangements as well as multiple adjacent divisions.

**Figure 9:**
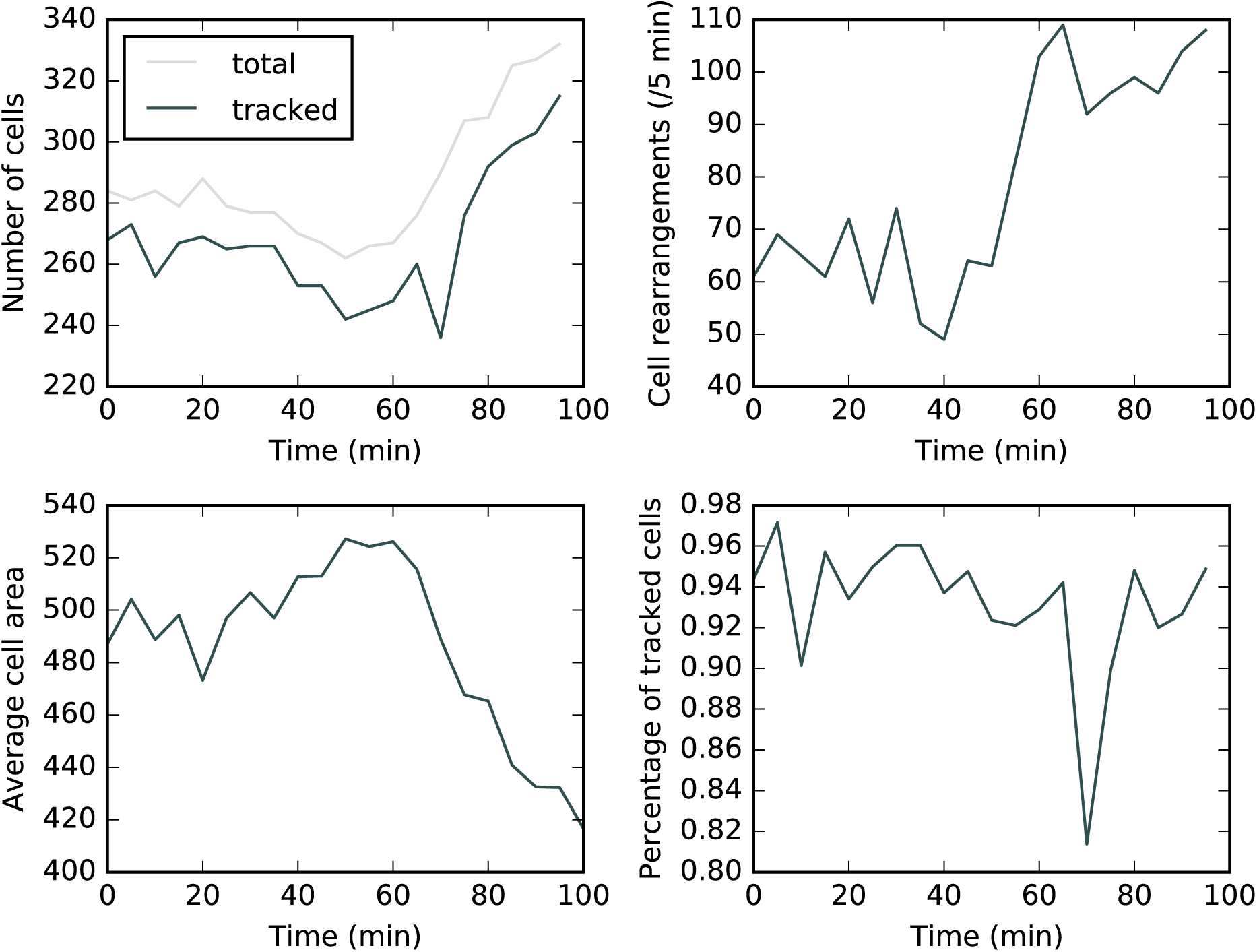
Tracking statistics of the *in vivo* dataset. (Top left) The total number of cells at each time point is constant initially and increases from 60 min onwards. The total number of tracked cells correlates with the total number of cells in the tissue. (Top right) The total number of rearrangements between successive time frames is measured by our algorithm. We record the total number of rearrangements as the total number of cells that either gain or loose neighbours between successive frames. (Bottom left) The average cell area in each frame decreases as the total number of cells increases. (Bottom right) The percentage of tracked cells decreases at around 70 min, when the amounts of cell rearrangement and division are highest.

Since cell rearrangements are one of the most difficult aspects of cell tracking, and our *in vivo* data exhibit a high frequency of such events, it is natural to ask what percentage of cells are correctly tracked. To estimate this percentage, we compare the results in figure 9, where up to 30% of cells are involved in neighbour exchange between frames in a population of up to 340 cells, with the results shown in figure 7, where in the case of 400 cells and 4% of edges undergoing T1 transitions between frames (corresponding to 30% of cells involved in neighbour exchanges) we find the percentage of correctly tracked cells to be 85%. This provides a lower bound for the success rate of the algorithm on the *in vivo* frames. When up to 3% of edges undergo T1 transitions (corresponding to 25% of cells in the tissue involved in neighbour exchanges), the success rate of the algorithm is 98%.

The tracking of epithelial *in vivo* data enables quantitative assessment of dynamic changes in cellular morphology. The tracking results in figure 9 reveals that the analysed section of the epidermis undergoes 60 cell rearrangements per five minutes initially and around 100 cell rearrangements per five minutes at the end of the observed time interval. The average ratio of the maximal area and the minimal area observed for individual cells during the period of measurement is 4.2, indicating that on average cells increase their apical area by a factor of four during mitotic rounding. A total of 18 cell deaths are tracked in the dataset. A striking feature is the level of T1 transitions occurring during this stage of development, even in the absence of gross morphogenetic movements found in earlier or later stages.

### Calculation times

To analyse the scaling of the calculation times with tissue size we repeat the permutation test with tissues of square dimension of varying size on a desktop computer with an Intel i5-6500T CPU (2.5GHz) and 8GB RAM. We find that the calculation times scale subquadratically with cell number (figure 10).

**Figure 10:**
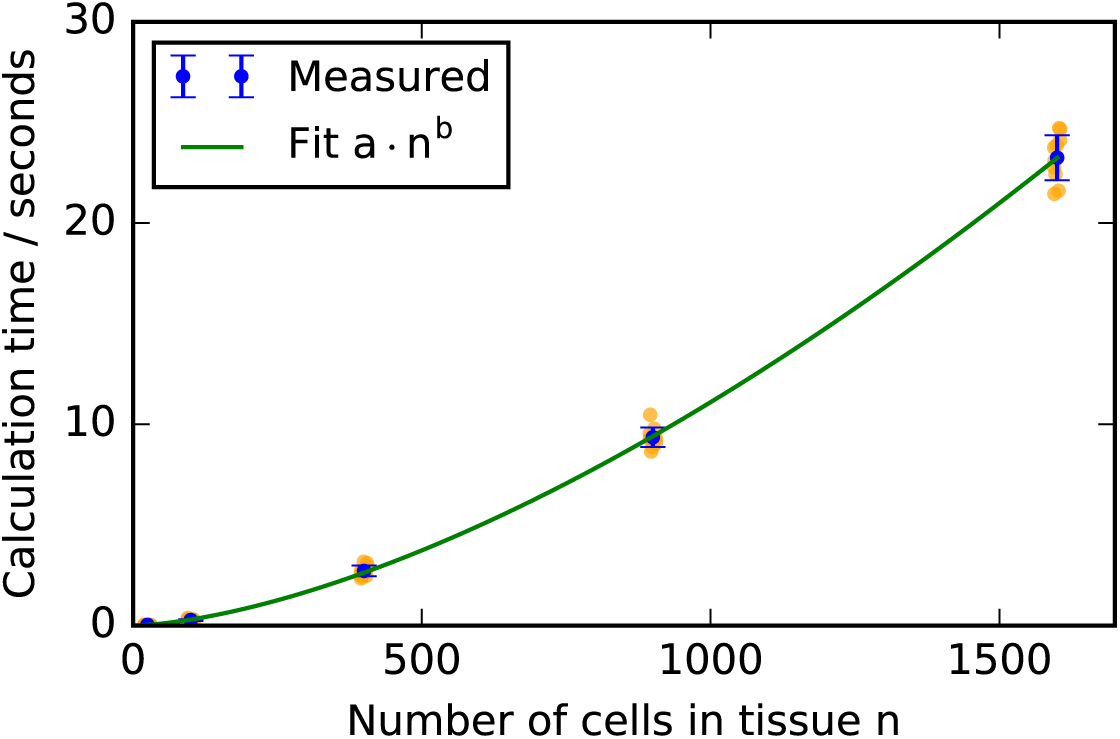
Scaling of the calculation times with tissue size. Square virtual tissues of varying sizes were generated and the calculation times of the algorithm under the permutation test in figure 6A recorded. Orange dots represent calculation times for individual realisations of the test and error bars denote the standard deviation. The exponent *b* of the polynomial fit is 1.6. The calculation times were measured on a desktop computer with an Intel i5-6500T CPU (2.5GHz) and 8GB RAM.

The calculation times for the experimental images analysed in figure 8 vary more widely than for the *in silico* datasets. For the tracking between the first and second frames in figure 8, the algorithm required 43 seconds to run, whereas between the second and the third frames the algorithm required 9 seconds. This is due to differences in the time required to find the first correct mapping; in the first example 154 cells were searched before the first correct mapping was found, whereas in the second example only 12 cells were searched. This means that the number of cells considered when finding the initial mappings depends on the graph structure of the analysed frames and impacts on the calculation time of the algorithm. In total, analysing all 21 frames of the *in vivo* data presented in figures 8 and 9 requires 19 minutes of calculation time.

## 4 Discussion

Cell tracking in epithelial sheets has the potential to generate a vast amount of quantitative data to inform our understanding of the contributions of different cellular processes to tissue morphogenesis. However, cell tracking is notoriously difficult, especially for the complex morphogenetic processes that occur as embryogenesis proceeds. Here, we present an algorithm based on MCS detection for the tracking of cells in segmented images of epithelial sheets. Our algorithm successfully tracks cells in *in vivo* images of the *Drosophila* embryonic epidermis, a challenging dataset compared to other tissues, as well as in randomly generated *in silico* datasets, without the need for the adjustment of tissue specific parameters such as weights for individual terms in a global minimisation scheme [14]. The use of *in silico* data to test our algorithm allows us to analyse the performance of our algorithm for a large range of experimentally observed cell rearrangements and tessellations.

The tracking of cells in *in vivo* datasets such as presented in figures 8 and 9 provides quantitative insight into tissue homeostasis. Using our algorithm we measure example quantities that would not be accessible without a robust cell tracking method. The amount of cell re-arrangement, the extent of mitotic rounding, and the occurrence of cell death in the observed frames each can be used to learn about tissue homeostasis in developing epithelia. Within the analysed dataset, we find a significant number of T1 transitions despite the absence of gross morphogenetic movements. This may be driven by the large number of proliferation events that occur. Further, using *in vivo* imaging together with our tracking algorithm allows the observation of cell death or cell delamination without the need for fluorescent markers of apoptosis. Future applications of the algorithm to such processes may, for example, provide novel insight to tissue size control in the *Drosophila* embryonic epidermis [9, 45] and can also be adapted to study the dynamics of epithelial wound closure. In this and other systems cell tracking may enable the observation of cell death due to delamination as opposed to apoptosis [46]. Access to quantification of cell rearrangement and area changes has recently provided insight to wing morphogenesis in *Drosophila* [43].

Our algorithm is able to track cells that undergo significant movement and neighbour exchanges between frames. For example, we can correctly track cells in tissues where more than 40% of the cells rearrange between successive movie frames (figure 7). In addition, even comparably large gaps in the initial MCS can be filled in during the post-processing step (figures 2 and 8). For example, in the first tracking step in figure 1, only 182 of the 246 tracked cells were identified by the MCS algorithm, and it was possible to track the 64 remaining cells during the post-processing step. For comparison, Heller et al [13] report 15 cell rearrangements per 1000 cells per hour at an imaging interval of six minutes for their time-lapse microscopy data of *Drosophila* wing imaginal discs. In addition, the experimental data shown in figures 2, 8, and 9 include junctions shared by four or more cells (yellow asterisks in figure 9) while our *in silico* data include multiple instances of such junctions (figure 6D). Therefore, higher-order junctions, such as multicellular rosettes [47, 48], do not pose a challenge to our algorithm.

Our algorithm is able to correctly track cells in all considered test cases. However, on rare occasions a few cells at the tissue boundary cannot be tracked. It may be possible to adapt the algorithm to track these cells, if this is considered necessary for the application at hand. In the current version of the algorithm, two connections to already tracked cells that are preserved between two time frames are a condition to add a cell-to-cell mapping in the post-processing algorithm. Further analysis of cases where this condition is not fulfilled may reveal ways to relax it.

When generating *in silico* data to test the algorithm, we used Voronoi tessellations in combination with Lloyd’s relaxation to generate data that resembles tissues in the *Drosophila* wing imaginal disc [33]. We expect the algorithm to perform less well on tissues whose network structure is nearly homogeneous. For example, in an epithelial sheet where cells are arranged in a hexagonal fashion, such as the early *Drosophila* embryonic epidermis [49] or the late pupal *Drosophila* wing [50], the local adjacency network of each cell is identical, and hence a network-based tracking algorithm may not be able to distinguish cells. When generating *in silico* tissues, we use four Lloyd’s relaxation steps after Voronoi tessellation. With each Lloyd’s relaxation step, the homogeneity of the tissue increases. We were able to successfully repeat all *in silico* tests on virtual tissues that were generated using up to 14 Lloyd’s relaxation steps. Hence, we expect the algorithm to be suitable for tissues that can be well described with 14 or fewer Lloyd’s relaxation steps, such as the chick neural tube embryonic epithelium, or the *Drosophila* eye disc [33].

The algorithm relies on being able to generate polygonal tessellations from segmented video microscopy data. In particular, all *in silico* tests we conducted consider tissues where each cell has at least three neighbours. Conceptually, it would be possible to apply the algorithm to tissues in which individual cells may have only two neighbours, although such examples have not been included in the present analysis.

In microscopy videos including division events we expect the algorithm to perform well in tissues in which no adjacent divisions occur between successive movie frames, and in which cells adjacent to the dividing cell do not undergo rearrangements before the next frame is captured. Our algorithm is designed to identify mother and daughter cells of a division event by establishing the bordering cells that gain an edge during the division event. In the case of two adjacent divisions, and if cells adjacent to a division event gain edges due to cell rearrangements, the dividing cell cannot be correctly identified. An example of a typical tracking error for two adjacent divisions is shown in figure 11. In cases where the division resolution step fails, our Python implementation returns all tracked cells of the post-processing step, and gives a warning that the division has not been resolved. In these cases, manual correction methods could be used for incorrectly tracked cells in the vicinity of division events.

**Figure 11:**
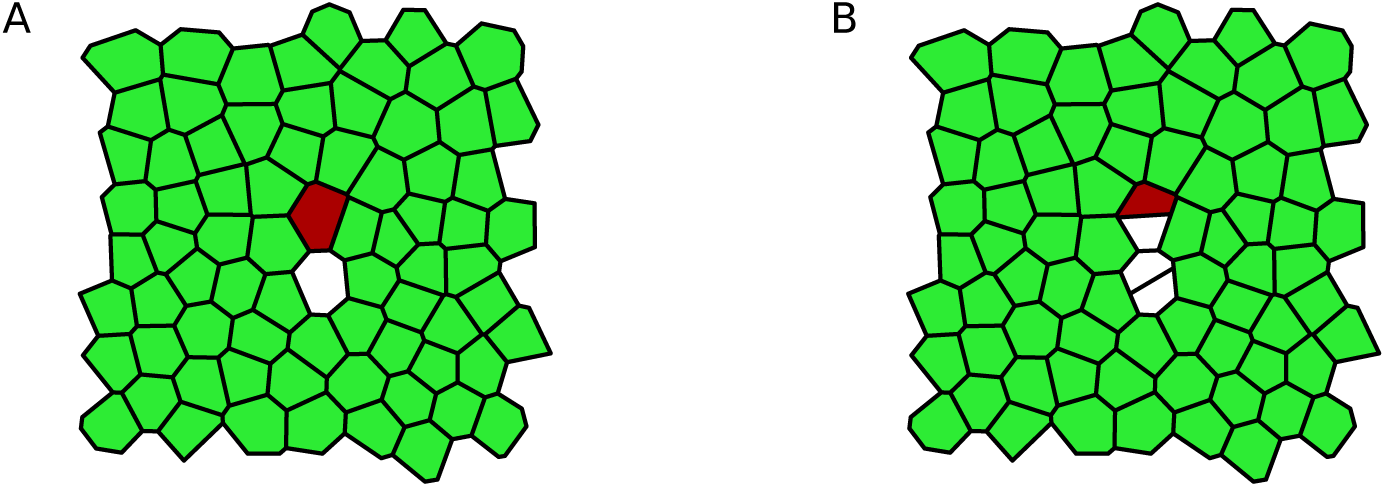
Tracking errors can occur if adjacent cells divide. Here, all green (light) cells are tracked correctly. One of the mother cells (red/dark) of the division events has been incorrectly associated with one of the daughter cells of the division.

The parameters of the algorithm are chosen to maximise its robustness and avoid the necessity to adjust the parameters to individual applications. For example, the cutoff length *d*_max_, that determines the distance below which two cells in consecutive movie frames are considered mappable to each other, was chosen to be 10 times the average cell length in the tissue, which is significantly larger than the movement that is to be expected between consecutive frames of a live-imaging microscopy video. However, parameter adjustments may be possible for individual applications in order to decrease the algorithm calculation times. For example, the size of the extended neighbourhood considered in the initial step or the iterative extension could be reduced to include only nearest neighbours instead of nearest neighbours and second nearest neighbours in case the tissue is sufficiently heterogeneous. Similarly, one might decrease the *d*_max_ for possible cell pairings if the cell positions are not expected to vary significantly between time frames.

Adjustments may be possible to extend the applicability of the algorithm to a wider range of tissues. For example, instead of automatic detection of the initial seeds for the MCS detection algorithm, a small set of seeds could be manually supplied to guide the tracking. This should improve the performance of the algorithm on homogeneous tissues. In such cases, irregular boundaries may also help to aid the initial seeding. During the construction of the MCS, non-adjacent cells are considered for addition to the MCS whenever the extension of the MCS by adjacent cells is not possible. An alternative option to extend an intermediate MCS may be to repeat the initial seeding algorithm.

In the present work, we have sought to keep geometrical input to the algorithm to a minimum. Cases where geometric data are taken into account comprise division events where one of the daughter cells is four- or three-sided, since in these cases we are not able to make a decision on which cell is the second daughter cell based on network adjacency alone. If future applications reveal cases where the algorithm performs poorly due to a large number of cell neighbour exchanges or high degree of tissue homogeneity, then it may be possible to construct algorithms that combine information on the network topology with data on cell shapes, cell positions and cell movements to improve performance. For example, information on network topology could be integrated into previous algorithms that minimise differences between geometric properties of cells, such as cell size and location [24], with information about network connectivity.

In cell tracking applications, the scaling of the algorithm with tissue size is crucial. Potential applications range from systems of 30 cells (*Drosophila* embryonic epidermal P compartments [9]), to 10,000 cells (*Drosophila* imaginal wing disc [31]; the wing pouch has about 3,000 cells [40]). Calculation times in the presented algorithm scale subquadratically with cell number, making it suitable for applications of varying sizes. For example, extrapolating the data in figure 10, a tissue of 10,000 cells could be tracked across two frames within 10 minutes. The scaling of the algorithm is polynomial despite the fact that it is based on MCS detection, which is known to scale exponentially in the general case. MCS detection has a wide range of research applications, including protein interaction networks [51, 52] and finding the binding sites of chemical structures [53]. Our approach of reducing the MCS search to a localised search may have applications in other areas where the networks are inherently planar.

Our algorithm is designed to track cells in segmented microscopy videos of epithelial sheets in two dimensions. However, it may be possible to apply the algorithm to datasets of epithelial sheets that are embedded in a three-dimensional environment, such as the *Drosophila* imaginal wing disc [4], or the *Drosophila* embryonic epidermis [6,9], including tissues that can be mapped onto a cylinder or ellipsoid, such as the mouse visceral endoderm [48].

A large number of cell tracking algorithms have been developed for various applications [10– 27]. Further efforts are required to compare these algorithms with our own, and to identify the algorithm best suited for an individual dataset. In the cell tracking challenge [54] the authors provide microscopy videos from a variety of *in vitro* cell cultures, including, for example, mouse embryonic stem cells and human squamous lung carcinoma cells, together with ground-truth segmentation and tracking data as benchmarks for cell tracking and segmentation algorithms. However, many of the published algorithms above have not yet been applied to the challenge, and benchmark datasets for epithelial sheets are not currently available. The fully segmented dataset published within the MCSTracker project can provide a benchmark for future epithelial cell tracking applications. In [55] *in silico* datasets are used as benchmarking datasets for particle tracking algorithms.

The proposed algorithm provides a two-dimensional tracking solution specialised for epithelial sheets that attempts to maximise the information that can be gained from the packing that is typical to such tissues. It may, however, be possible to extend this algorithm to applications of cell tracking where cells are not physically connected by constructing adjacency networks from Voronoi tessellations that use the cell locations as seeds. We hope that, as segmentation tools are developed further, the combination of our algorithm with these tools will lead to further insights into cellular behaviour in epithelial tissues.

## Acknowledgements

J.K. acknowledges funding from the Engineering and Physical Sciences Research Council through a studentship. J.Z. acknowledges funding support from the National Science Foundation (awards CBET-1553826 and CBET- 1403887). The authors thank Cody Narciso for sharing the mi- croscopy data.

## References

1. Stephens, D. J. and Allan, V. J. Light microscopy techniques for live cell imaging. Science, 300(5616):82–86, 2003. 10.1126/science.1082160.

2. Pantazis, P. and Supatto, W. Advances in whole-embryo imaging: a quantitative transition is underway. Nat. Rev. Mol. Cell. Biol., 15(5):327–339, 2014. 10.1038/nrm3786.

3. Truong, T. V. and Supatto, W. Toward high-content/high-throughput imaging and analysis of embryonic morphogenesis. Genesis, 49(7):555–569, 2011. 10.1002/dvg.20760.

4. Mao, Y., Tournier, A. L., Bates, P. A., Gale, J. E., Tapon, N., and Thompson, B. J. Planar polarization of the atypical myosin Dachs orients cell divisions in *Drosophila*. Genes Dev., 25(2):131–136, 2011. 10.1101/gad.610511.

5. Gibson, M. C., Patel, A. B., Nagpal, R., and Perrimon, N. The emergence of geometric order in proliferating metazoan epithelia. Nature, 442(7106):1038–1041, 2006. 10.1038/nature05014.

6. Rauzi, M., Verant, P., Lecuit, T., and Lenne, P.-F. Nature and anisotropy of cortical forces orienting *Drosophila* tissue morphogenesis. Nat. Cell Biol., 10(12):1401–1410, 2008. 10.1038/ncb1798.

7. Collinet, C., Rauzi, M., Lenne, P.-F., and Lecuit, T. Local and tissue-scale forces drive oriented junction growth during tissue extension. Nat. Cell Biol., 17(10):1247–1258, 2015. 10.1038/ncb3226.

8. Ritsma, L., Ellenbroek, S. I. J., Zomer, A., Snippert, H. J., de Sauvage, F. J., Simons, B. D., Clevers, H., and van Rheenen, J. Intestinal crypt homeostasis revealed at single-stem-cell level by *in vivo* live imaging. Nature, 507(7492):362–365, 2014. 10.1038/nature12972.

9. Parker, J. Control of compartment size by an EGF ligand from neighboring cells. Curr. Biol., 16(20):2058–2065, 2006. 10.1016/j.cub.2006.08.092.

10. Mashburn, D. N., Lynch, H. E., Ma, X., and Hutson, M. S. Enabling user-guided segmentation and tracking of surface-labeled cells in time-lapse image sets of living tissues. Cytometry A, 81A(5):409–418, 2012. 10.1002/cyto.a.22034.

11. Cilla, R., Mechery, V., Hernandez de Madrid, B., Del Signore, S., Dotu, I., and Hatini, V. Segmentation and tracking of adherens junctions in 3D for the analysis of epithelial tissue morphogenesis. PLoS Comput. Biol., 11(4):e1004124, 2015. 10.1371/journal.pcbi.1004124.

12. Schiegg, M., Hanslovsky, P., Kausler, B., Hufnagel, L., and Hamprecht, F. Conservation tracking. IEEE Int. Comp. Vis., 2928–2935, 2013. 10.1109/ICCV.2013.364.

13. Heller, D., Hoppe, A., Restrepo, S., Gatti, L., Tournier, A., Tapon, N., Basler, K., and Mao, Y. EpiTools: An open-source image analysis toolkit for quantifying epithelial growth dynamics. Dev. Cell, 36(1):103–116, 2016. 10.1016/j.devcel.2015.12.012.

14. Padfield, D., Rittscher, J., and Roysam, B. Coupled minimum-cost flow cell tracking for high-throughput quantitative analysis. Med. Image Anal., 15(4):650–668, 2011. 10.1016/j.media.2010.07.006.

15. Youssef, S., Gude, S., and Radler, J. O. Automated tracking in live-cell time-lapse movies. Integr. Biol., 3:1095–1101, 2011. 10.1039/C1IB00035G.

16. Wait, E., Winter, M., Bjornsson, C., Kokovay, E., Wang, Y., Goderie, S., Temple, S., and Cohen, A. Visualization and correction of automated segmentation, tracking and lineaging from 5-D stem cell image sequences. BMC Bioinform., 15(1):328, 2014. 10.1186/1471-2105- 15-328.

17. Winter, M., Wait, E., Roysam, B., Goderie, S. K., Ali, R. A. N., Kokovay, E., Temple, S., and Cohen, A. R. Vertebrate neural stem cell segmentation, tracking and lineaging with validation and editing. Nat. Protocols, 6(12):1942–1952, 2011. 10.1038/nprot.2011.422.

18. Sommer, C., Straehle, C., K¨othe, U., and Hamprecht, F. A. Ilastik: Interactive learning and segmentation toolkit. In IEEE International Symposium on Biomedical Imaging: From Nano to Macro, 230–233. 2011. 10.1109/ISBI.2011.5872394.

19. Liu, K., Lienkamp, S. S., Shindo, A., Wallingford, J. B., Walz, G., and Ronneberger, O. Optical flow guided cell segmentation and tracking in developing tissue. In IEEE 11th International Symposium on Biomedical Imaging (ISBI), 298–301. 2014. 10.1109/ISBI.2014.6867868.

20. Bellaiche, Y., Bosveld, F., Graner, F., Mikula, K., Remesikova, M., and Smisek, M. New robust algorithm for tracking cells in videos of *Drosophila* morphogenesis based on finding an ideal path in segmented spatio-temporal cellular structures. In IEEE Annu. Int. Conf. Eng. Med. Biol. Soc., 6609–6612. 2011. 10.1109/IEMBS.2011.6091630.

21. Aly, A. A., Deris, S. B., and Zaki, N. Intelligent algorithms for cell tracking and image segmentation. Int. J. Comput. Sci. Inf. Technol., 6(5):21–37, 2014.

22. Wang, Q., Niemi, J., Tan, C.-M., You, L., and West, M. Image segmentation and dynamic lineage analysis in single-cell fluorescence microscopy. Cytometry A, 77A(1):101–110, 2010. 10.1002/cyto.a.20812.

23. Raffel, M., Willert, C. E., Wereley, S., and Kompenhans, J. Particle Image Velocimetry: A Practical Guide. Springer-Verlag Berlin Heidelberg, 2007.

24. Puliafito, A., Hufnagel, L., Neveu, P., Streichan, S., Sigal, A., Fygenson, D. K., and Shraiman, B. I. Collective and single cell behavior in epithelial contact inhibition. Proc. Natl. Acad. Sci. U.S.A., 109(3):739–744, 2012. 10.1073/pnas.1007809109.

25. Al-Kofahi, Y., Lassoued, W., Lee, W., and Roysam, B. Improved automatic detection and segmentation of cell nuclei in histopathology images. IEEE Trans. Biomed. Eng., 57(4):841–852, 2010. 10.1109/TBME.2009.2035102.

26. Amat, F., Lemon, W., Mossing, D. P., McDole, K., Wan, Y., Branson, K., Myers, E. W., and Keller, P. J. Fast, accurate reconstruction of cell lineages from large-scale fluorescence microscopy data. Nat. Meth., 11(9):951–958, 2014. 10.1038/nmeth.3036.

27. Aigouy, B., Farhadifar, R., Staple, D. B., Sagner, A., Röper, J.-C., Jülicher, F., and Eaton, S. Cell flow reorients the axis of planar polarity in the wing epithelium of *Drosophila*. Cell, 142(5):773–786, 2010. 10.1016/j.cell.2010.07.042.

28. Hoebe, R. A., Van Oven, C. H., Gadella, T. W. J., Dhonukshe, P. B., Van Noorden, C. J. F., and Manders, E. M. M. Controlled light-exposure microscopy reduces photobleaching and phototoxicity in fluorescence live-cell imaging. Nat. Biotech., 25(2):249–253, 2007. 10.1038/nbt1278.

29. Wood, W. and Jacinto, A. Cell Migration: Developmental Methods and Protocols, chapter Imaging Cell Movement During Dorsal Closure in *Drosophila* Embryos, 203–210. Humana Press, Totowa, NJ, 2005. 10.1385/1-59259-860-9:203.

30. Mavrakis, M., Rikhy, R., Lilly, M., and Lippincott-Schwartz, J. Fluorescence Imaging Techniques for Studying *Drosophila* Embryo Development. John Wiley & Sons, Inc., 2001. 10.1002/0471143030.cb0418s39.

31. Farhadifar, R., Röper, J.-C., Aigouy, B., Eaton, S., and Jölicher, F. The influence of cell mechanics, cell-cell interactions, and proliferation on epithelial packing. Curr. Biol., 17(24):2095–2104, 2007. 10.1016/j.cub.2007.11.049.

32. Escudero, L. M., da F. Costa, L., Kicheva, A., Briscoe, J., Freeman, M., and Babu, M. M. Epithelial organisation revealed by a network of cellular contacts. Nat. Commun., 2:526, 2011. 10.1038/ncomms1536.

33. Sánchez-Gutiérrez, D., Tozluoglu, M., Barry, J. D., Pascual, A., Mao, Y., and Escudero, L. M. Fundamental physical cellular constraints drive self-organization of tissues. EMBO J., 2015. 10.15252/embj.201592374.

34. Sáez, A., Acha, B., Montero-Sánchez, A., Rivas, E., Escudero, L. M., and Serrano, C. Neuromuscular disease classification system. J. Biomed. Opt., 18(6):066017–066017, 2013. 10.1117/1.JBO.18.6.066017.

35. Ullmann, J. R. An algorithm for subgraph isomorphism. J. ACM, 23(1):31–42, 1976. 10.1145/321921.321925.

36. Krissinel, E. B. and Henrick, K. Common subgraph isomorphism detection by backtracking search. Software Pract. Exper., 34(6):591–607, 2004. 10.1002/spe.588.

37. Osborne, J. M., Bernabeu, M. O., Bruna, M., Calderhead, B., Cooper, J., Dalchau, N., Dunn, S.-J., Fletcher, A. G., Freeman, R., Groen, D., et al. Ten simple rules for effective computational research. PLoS Comput. Biol., 10(3):e1003506, 2014. 10.1371/journal.pcbi.1003506.

38. Hagberg, A. A., Schult, D. A., and Swart, P. J. Exploring network structure, dynamics, and function using NetworkX. In Proceedings of the 7th Python in Science Conference (SciPy2008), 11–15. Pasadena, CA USA, 2008.

39. Honda, H. Description of cellular patterns by Dirichlet domains: The two-dimensional case. J. Theor. Biol., 72(3):523–543, 1978. 10.1016/0022-5193(78)90315-6.

40. Narciso, C., Wu, Q., Brodskiy, P. A., Garston, G., Baker, R. E., Fletcher, A. G., and Zartman, J. J. Patterning of wound-induced intercellular Ca 2+ flashes in a developing epithelium. Phys. Biol., 12(5):056005, 2015. 10.1088/1478-3975/12/5/056005.

41. Parton, R. M., Vallés, A. M., Dobbie, I. M., and Davis, I. Collection and mounting of *Drosophila* embryos for imaging. Cold Spring Harbor Protocols, 2010(4):pdb.prot5403, 2010. 10.1101/pdb.prot5403.

42. Nagai, T., Kawasaki, K., and Nakamura, K. Vertex dynamics of two-dimensional cellular patterns. J. Phys. Soc. Jpn., 57(7):2221–2224, 1988. 10.1143/JPSJ.57.2221.

43. Etournay, R., Popović, M., Merkel, M., Nandi, A., Blasse, C., Aigouy, B., Brandl, H., Myers, G., Salbreux, G., Jülicher, F., et al. Interplay of cell dynamics and epithelial tension during morphogenesis of the *Drosophila* pupal wing. eLife, 4:e07090, 2015. 10.7554/eLife.07090.

44. Aegerter-Wilmsen, T., Smith, A. C., Christen, A. J., Aegerter, C. M., Hafen, E., and Basler, K. Exploring the effects of mechanical feedback on epithelial topology. Development, 137(3):499–506, 2010. 10.1242/dev.041731.

45. Kursawe, J., Brodskiy, P. A., Zartman, J. J., Baker, R. E., and Fletcher, A. G. Capabilities and limitations of tissue size control through passive mechanical forces. PLoS Comput. Biol., 11(12):1–26, 2016. 10.1371/journal.pcbi.1004679.

46. Marinari, E., Mehonic, A., Curran, S., Gale, J., Duke, T., and Baum, B. Live-cell delamination counterbalances epithelial growth to limit tissue overcrowding. Nature, 484(7395):542–545, 2012. 10.1038/nature10984.

47. Blankenship, J. T., Backovic, S. T., Sanny, J. S. P., Weitz, O., and Zallen, J. A. Multicellular Rosette Formation Links Planar Cell Polarity to Tissue Morphogenesis. Developmental Cell, 11(4):459–470, 2006. 10.1016/j.devcel.2006.09.007.

48. Trichas, G., Smith, A. M., White, N., Wilkins, V., Watanabe, T., Moore, A., Joyce, B., Sugnaseelan, J., Rodriguez, T. A., Kay, D., et al. Multi-cellular rosettes in the mouse visceral endoderm facilitate the ordered migration of anterior visceral endoderm cells. PLoS Biol., 10(2):e1001256, 2012. 10.1371/journal.pbio.1001256.

49. Warn, R. and Magrath, R. F-actin distribution during the cellularization of the drosophila embryo visualized with FL-phalloidin. Exp. Cell Res., 143(1):103–114, 1983. http://dx.doi.org/10.1016/0014-4827(83)90113-1.

50. Classen, A.-K., Anderson, K. I., Marois, E., and Eaton, S. Hexagonal packing of *Drosophila* wing epithelial cells by the planar cell polarity pathway. Dev. Cell, 9(6):805–817, 2005. 10.1016/j.devcel.2005.10.016.

51. Ciriello, G., Mina, M., Guzzi, P. H., Cannataro, M., and Guerra, C. Alignnemo: A local network alignment method to integrate homology and topology. PLoS ONE, 7(6):e38107, 2012. 10.1371/journal.pone.0038107.

52. Aladağ, A. E. and Erten, C. Spinal: scalable protein interaction network alignment. Bioinformatics, 29(7):917–924, 2013. 10.1093/bioinformatics/btt071.

53. Raymond, J. W. and Willett, P. Maximum common subgraph isomorphism algorithms for the matching of chemical structures. J. Comput.-Aided Mol. Des., 16(7):521–533, 2002. 10.1023/A:1021271615909.

54. Maška, M., Ulman, V., Svoboda, D., Matula, P., Matula, P., Ederra, C., Urbiola, A., España, T., Venkatesan, S., Balak, D. M., et al. A benchmark for comparison of cell tracking algorithms. Bioinformatics, 30(11):1609–1617, 2014. 10.1093/bioinformatics/btu080.

55. Chenouard, N., Smal, I., de Chaumont, F., Maska, M., Sbalzarini, I. F., Gong, Y., Car-dinale, J., Carthel, C., Coraluppi, S., Winter, M., et al. Objective comparison of particle tracking methods. Nat. Meth., 11(3):281–289, 2014. 10.1038/nmeth.2808.

